# Tunable multivalent Fe(II)-based glycoassemblies as mimetics for native high-mannose glycans

**DOI:** 10.64898/2025.12.17.694979

**Authors:** Emerson Hall, Yu-Shien Sung, Chad W. Priest, Julia M. Stauber, Alex J. Guseman

## Abstract

High Mannose Glycans (HMGs) play key roles in eukaryotic biology, regulating processes ranging from protein folding to host pathogen defense. Lectins have evolved to interact with these glycans through multivalent interactions facilitated by the multiple sugars displayed on glycans and via multiple binding sites on each lectin. Using Fe(II) iminopyridine complexes, we generated chemically defined multivalent glycan displays where the valency, arm length, and spatial display of mannose residues can be controlled via subcomponent synthesis. Due to its sensitivity towards the geometric display of mannose residues, monomeric Griffithsin (mGRFT) was utilized as a model lectin. Interactions between the Fe(II) glycan assemblies and mGRFT were characterized using biolayer interferometry (BLI), isothermal titration calorimetry (ITC), and NMR spectroscopy. Our results display a >1000-fold range in K_D_ for Fe(II) iminopyridine complexes that can be tuned by factors such as saccharide tether length and number of sugars displayed. Through leveraging systematic molecular-level modifications, we demonstrate that tunable Fe(II) glycan assemblies can be used both as mimetics for high mannose glycans as well as competitive inhibitors for native glycan binding.

## Introduction

N-linked glycosylation is a highly conserved and essential post-translational modification in which oligosaccharides are covalently attached to the amide side chains of asparagine residues on target proteins.^1^ Dysregulation of N-glycans is implicated in a wide range of diseases, including congenital disorders, cancers, and viral infections, largely due to the influential role of N-glycans in physiological processes such as immune function, cell-cell communication, and host-pathogen interactions.^2–4^ N-linked glycans are classified into three main subtypes: high-mannose, hybrid, and complex, which each play critical roles in protein folding,^5^ stability, trafficking, and molecular recognition.^2^ Despite this diversity, all N-glycans share a conserved Man_3_GlcNAc_2_ core with variations in mannose branching and extension that generate multivalent displays of glycan epitopes which mediate avidity-driven lectin recognition across wide-ranging biological systems.

Lectins, which typically bind individual monosaccharides weakly (mM *K*_d_ range)^6^, achieve markedly higher affinity when engaging these multivalent glycan displays (μM–nM *K*_d_ range)^7,8^. Such multivalency can arise from the presence of multiple binding sites within a single lectin domain or from the cooperative activity of multiple lectin domains within an oligomeric assembly.^9^ The branched, antennary structure of glycans further enhances this effect, as lectins have evolved to recognize specific branch points or to interact with sugars presented in defined spatial geometries on glycan antennary arms. Although lectin specificity and multivalent binding are central to many biological processes, the precise structural factors that govern these interactions remain difficult to define, motivating the use of synthetic mimics to probe how glycan architecture directs lectin recognition.^8^ However, the intrinsic branching and dense spatial clustering that enable selective recognition of N-linked glycans make such systems difficult to replicate synthetically. Due to these complex structures and the difficultly to systematically modify them, achieving molecular-level control in the design of well-defined glycomimetics capable of probing glycan-based multivalent interactions remains a major challenge.^10–12^

**Scheme 1:**
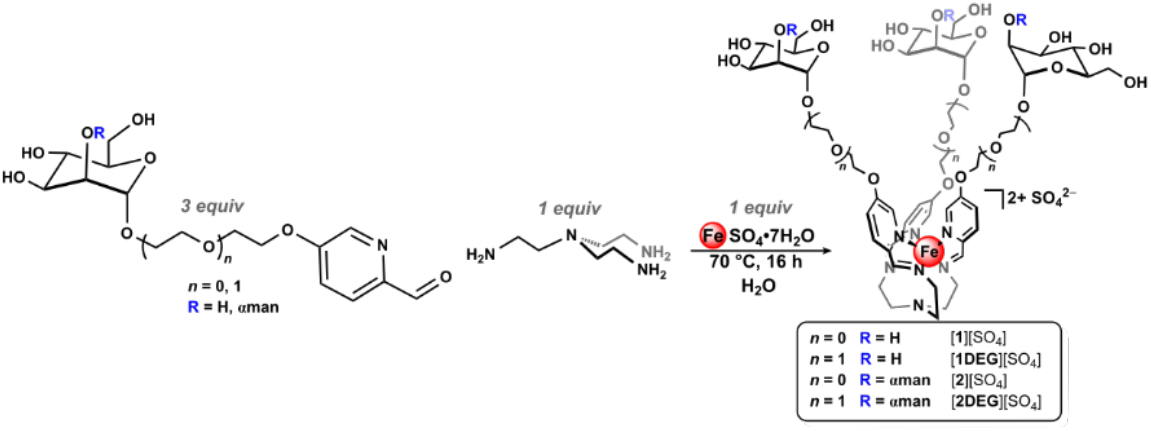
One pot reaction for the generation of metalloglycan architectures

To address this synthetic gap, we have recently introduced a modular platform based on coordination-driven subcomponent self-assembly,^13–15^ which employs metal ion templates to generate complex metalloglycan architectures from simple precursors in one pot.^16–18^ This platform integrates spatial precision, architectural tunability, and surface modularity to enable molecular-level control over glycan valency on well-defined three-dimensional frameworks. Unlike traditional polymeric glycomaterials,^19–21^ dendrimers,^22,23^ or nanoparticles,^24,25^ which often lack atomic-level definition and are difficult to systematically and precisely modify, our assemblies offer a unique combination of well-defined symmetry, tunable architecture, and modular functionalization through a synthetic “mix-and-match” molecular strategy. In this process, formyl-pyridine building blocks bearing glycan units are paired with multitopic amines to spontaneously self-assemble around iron(II) templates through the formation of covalent C=N linkages and dative N→Fe coordination bonds (Scheme 1).^26,27^ This self-assembly reaction provides rapid access to air- and water-stable Fe(II) iminopyridine complexes with well-defined carbohydrate arrangements that exhibit excellent stability under physiological conditions and demonstrate lectin-binding affinities in the micromolar to nanomolar range.^16–18^ Using this approach, we previously prepared well-defined Fe-based glycomimetics bearing between three and seventy-two structurally-defined appended saccharides. We have also shown that modulating the polyethylene glycol (PEG) linker length connecting the saccharide unit to the formylpyridine subcomponent offers a simple strategy to tune the glycan spatial and topological arrangement around the Fe(II) core.^17^ Modulating tether length allows us to adjust saccharide spacing and flexibility to enable controlled intersaccharide separation while providing opportunities to tailor molecular architectures to the specific binding surfaces of lectin targets. In this work, we leverage this precise control to systematically probe protein-glycan interactions by modulating molecular-level factors that govern multivalent binding.

As a model for probing multivalent oligomannose interactions, we selected the mannose-binding protein griffithsin (GRFT), a lectin isolated from the genus of red alga *Griffithsia*.^28^ GRFT, is a homodimeric protein with three mannose-binding sites per subunit that has been extensively characterized biophysically and structurally due to its potent antiviral activity, making it an ideal model system for testing glycomimetics.^28–33^ GRFT is a domain swapped dimer, consisting of two 121 amino acids polypetides that strand exchange to form beta sandwich structures, which recognize the terminal mannoses on high mannose glycans (HMGs) *via* a symmetric tripartite binding site (Fig. 1C).^34^ Insertion of additional glycine and serine residues in the hinge region of GRFT results in the formation of a stable monomer (mGRFT) with a single tripartite binding site.^33^ Binding of GRFT to free mannose,^33,35,36^ linkage specific mannose disaccharides,^33,35^ Man-9 glycan,^33,37^ and a variety of multivalent mannose displays^38^ have demonstrated that GRFT interacts with the α-1,2 linked terminal mannose residues attached on the two terminal mannose branches of a single N-glycan. X-ray crystallography studies reveal that the terminal mannose on the third branch of Man-9 cannot reach the third binding site of GRFT, which supports the hypothesis that crosslinking of glycoproteins occurs when this third binding site is occupied by a mannose residue from a second glycan.^33^

**Figure 1:**
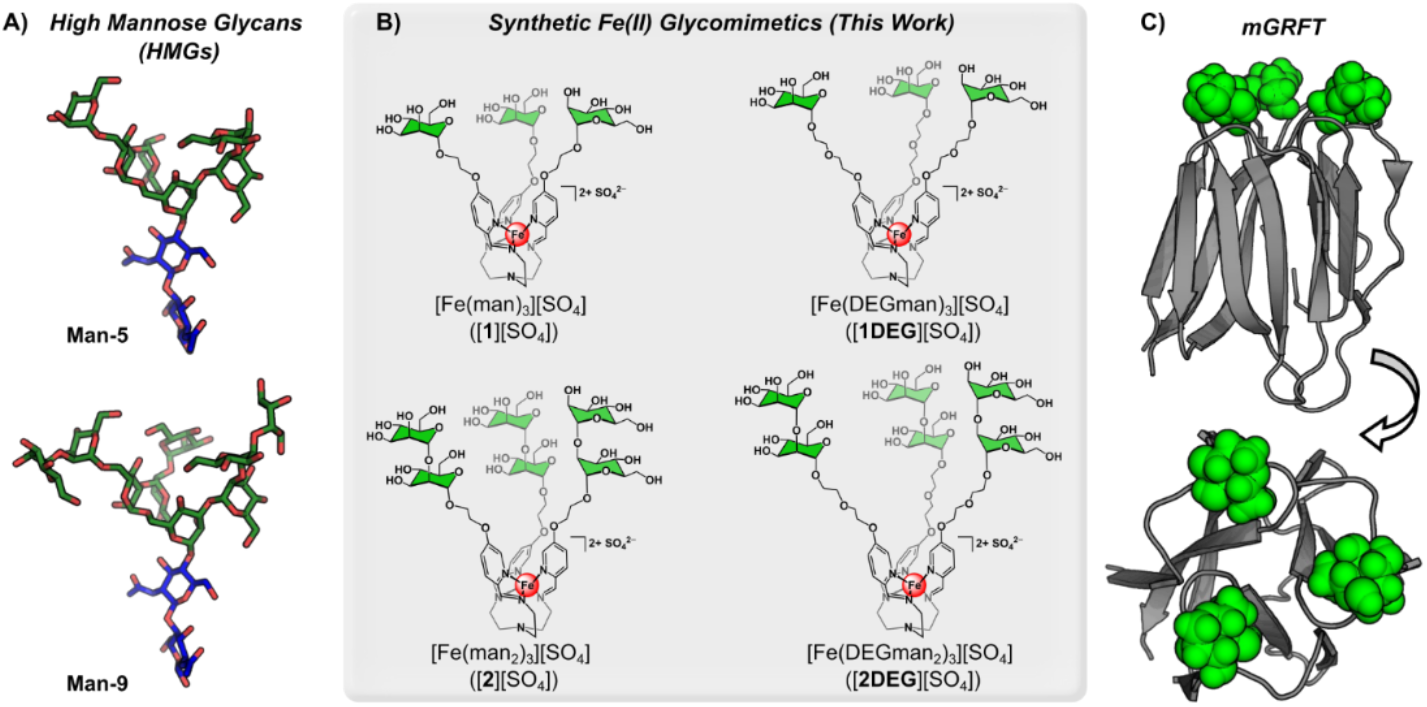
Glycan, glycomimetic, and mGRFT structures: A) Man-5 and Man-9 energy minimized structures generated via glycam and visualized in PYMOL using SNFG color coding (blue: N-acetyl glucosamine, green: mannose). B) Structures of Fe(II) glycan complexes with mannose units highlighted in green. C) Ribbon structure of mGRFT in grey (PDB: 2guc, 3ll2) with mannose groups as green spheres visualized in PYMOL.

Here, we investigate a series of multivalent Fe(II)-based glycoassemblies to mimic the branched mannose presentation of native N-linked glycans by characterizing their binding with mGRFT using nuclear magnetic resonance (NMR) spectroscopy, biolayer interferometry (BLI), and isothermal titration calorimetry (ITC). These results demonstrated that the parent trimannose Fe(II) iminopyridine complexes (Fig. 1B) mimic the tri-antennary branching structure of high mannose glycans and display mannose residues similarly to Man-5 and Man-6 glycans (Fig. 1A). We then examine the extensions of the molecular tether arms through the addition of ethylene glycol (EG) linkers or the display of mannose disaccharides to show binding behavior that mimics Man-9 glycans. Furthermore, we demonstrate that the ability to tune both linker length and local mannose valency can be applied combinatorically to rationally design a competitive inhibitor that surpasses the affinity of mGRFT for its native ligand, Man-9.

## Results and Discussion

### Synthesis of Fe(II)-based glycomimetics

We began these studies using our previously reported Fe(II)-anchored trimannose complex, [**1**][SO_4_],^16^ which is prepared through the condensation of 5-substituted ethyloxymannose formylpyridine with tris-2aminoethylamine (tren) in the presence of FeSO_4_•7H_2_O. This reaction yields a trivalent Fe(II)-anchored glycan assembly that features a facial saccharide display, as shown in Fig. 1B. We envisioned this glycan geometry could complement the binding surface of GRFT, which features three equidistant mannose-binding sites per subunit that are oriented in a triangular arrangement.^39^ Given the tunable nature of this synthetic platform, we also employed our previously reported diethylene glycol-extended mannose formylpyridine variant,^17^ in addition to α-1,2-mannose mono- and diethylene glycol-linked picolinaldehyde subcomponents that we introduce in this work. These new organic building blocks were fully characterized by ^1^H and ^13^C NMR spectroscopy, IR spectroscopy, and high-resolution electrospray mass spectrometry (HR-ESIMS(+)).

Using this series of subcomponents, we prepared the corresponding iminopyridine Fe(II) complexes under the same reaction conditions established for the preparation of [**1**][SO_4_] to introduce three new complexes that systematically vary in tether length (diethyleneglycol extended derivative: [**1DEG**][SO_4_]) and the number of terminal mannose units (α-1,2-mannose derivatives: [**2**][SO_4_], and [**2DEG**][SO_4_], Fig. 1B). This series extends beyond our original trivalent Fe(II) glycoassembly to better mimic natural oligomannoside structures and to probe GRFT interactions with greater precision. All three new Fe(II) glycomimetics were characterized by ^1^H NMR spectroscopy, IR spectroscopy, UV-vis spectroscopy, and HR-ESIMS(+).

### [1][SO_4_] as a mimetic for Man-5 and Man-6 glycans

We first evaluated the binding of [**1**]^2+^ and Man-5 to mGRFT as initial reference points using BLI. Through kinetic fitting and steady state analysis, mGRFT displayed a mid-micromolar affinity towards Man-5, K_D_ = 40 ± 10 µM, respectively (Fig. 2A). These measurements align with the affinity of 3α,6α Mannopentaose (the core structure of Man-5) as determined through a series of 2D ^1^H-^15^N HSQC NMR titrations (K_D_ = 80 ± 10 µM). In comparison, the dissociation constant for the binding of mGRFT with [**1**]^2+^ was calculated as K_D_ = 1.09 ± 0.08 μM (Fig. 2B). This interaction is roughly comparable that of mGRFT with Man-5 and 3α,6α Mannopentaose, yet two orders of magnitude stronger than GRFT’s affinity towards free mannose (K_D_ = 102 μM),^35^ which is a result largely driven by multivalent effects. To further interrogate this interaction, we performed ITC binding studies with mGRFT and [**1**]^2+^. The data were fit to a one-site binding model, which resulted in an average *K*_d_ value of 13.37 ± 3.7 μM (Fig. 2C), and a binding stoichiometry (*n*) of 1.7. These data are consistent with a model in which a single [**1**]^2+^ molecule engages two of the three mGRFT binding sites but due to size constraints, the single complex is unable to fully engage all three sites. Supporting this interpretation, molecular docking studies conducted in Molecular Operating Environment (MOE)^40^ show that [**1**]^2+^ cannot simultaneously contact all three binding site. Fig 2F presents three views of this docking model, highlighting how the arms of [**1**]^2+^ are unable to fully reach the third site depicted in orange. An alternative model is that one [**1**]^2+^ molecule forms strong interactions at two sites while simultaneously establishing a weaker, transient interaction with the third site due to size limitations that prevent full engagement of all three binding locations.

**Figure 2:**
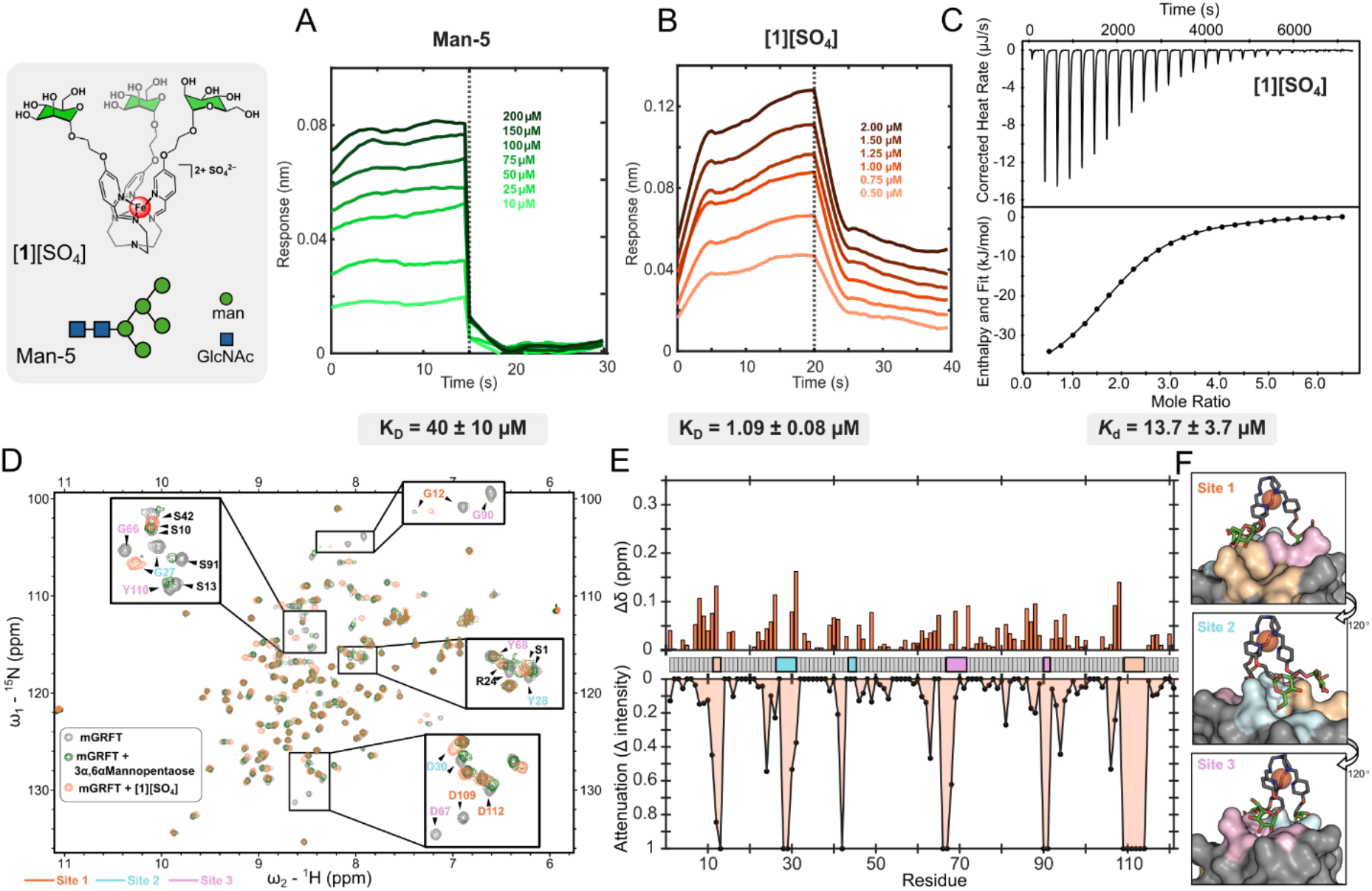
A) BLI trace with biotinylated Man-5 as the ligand with varying concentration of mGRFT as the analyte. B) BLI trace with mGRFT as the ligand with varying concentration of [**1**]^2+^ as the analyte. C) ITC thermograms (top) and fitted binding isotherms (bottom) for the binding of [**1**]^2+^ and mGRFT. D) 2D ^1^H-^15^N HSQC NMR spectrum of mGRFT alone (grey) and mGRFT bound with [**1**]^2+^ (coral) or 3α,6α Mannopentaose (green). E) Chemical shift perturbations, top, and changes in signal intensity, bottom, as a function of amino acid residue. F) MOE docking of [**1**]^2+^ onto mGRFT with the molecular surface colored based on binding site.

To understand the structural aspects governing the binding of [**1**]^2+^ to mGRFT, 2D ^1^H-^15^N HSQC spectra of the protein ligand complex were collected and analyzed for chemical shift perturbations and attenuations. These data revealed similarities in the chemical shift perturbation and attenuation profiles when compared with those of mGRFT bound to the core mannoses of Man-5 (3α,6α Mannopentaose) (Fig. 2D). Most dramatically, the signals attributed to the aspartic acid residues D67 (site 3) and D109 (site 1) of mGRFT fully attenuate when bound to [**1**]^2+^ (Fig. 2E), aligning with literature that reports full signal attenuation of the same residues when bound to a linear glycomimetic peptide,^38^ compared to a slight chemical shift when weakly bound to free mannose.^36^ These attenuations, compared to chemical shift perturbations (CSPs), suggest a weaker and more transient interaction with multiple bound conformations sampled rather than formation of a stable complex. These data are consistent with the structural model for mGRFT, in which three mannose binding sites are nearly identical in sequence and affinity. Consequently, the NMR spectra of bound mimetics reflect averaged signals of a heterogeneous mixture of bound states. These data suggest that the Fe(II) iminopyridine core plays a similar role to Man-3 branching core found in high mannose glycans to organize the appended mannose residues into a geometry that mirrors their presentation in Man-5 and Man-6 glycans. As a result, the [**1**]^2+^ scaffold serves as a functional glycomimetic of these high-mannose motifs.

### Extension of [1][SO_4_] arms mimics Man-9 binding

Combined insights from NMR, ITC, BLI, and docking studies suggest the mannose arrangement in [**1**]^2+^ restricts its full engagement with mGRFT, which highlights an opportunity to exploit molecular design principles to enable the synthesis of a glycomimetic that more closely resembles the binding strength of the native Man-9 ligand (67 ± 1 nM, Fig. 3A). Leveraging the synthetic tunability of this platform, we pursued two complementary strategies to probe how both antennary arm distance and saccharide topology influence lectin recognition. To examine distance effects, we first prepared the [Fe(DEGman)_3_][SO_4_] ([**1DEG**][SO_4_], Fig. 1B) analog, which introduces a single additional ethylene glycol (EG) spacer to extend the mannose tether. The added EG unit increases both the reach and conformational flexibility of arm, which we envisioned would better position the displayed mannose residues to access mGRFT’s third binding site. This extension was designed to more closely mimic the architecture of HMGs such as Man-9, and enhance the binding affinity into the nanomolar regime

**Figure 3:**
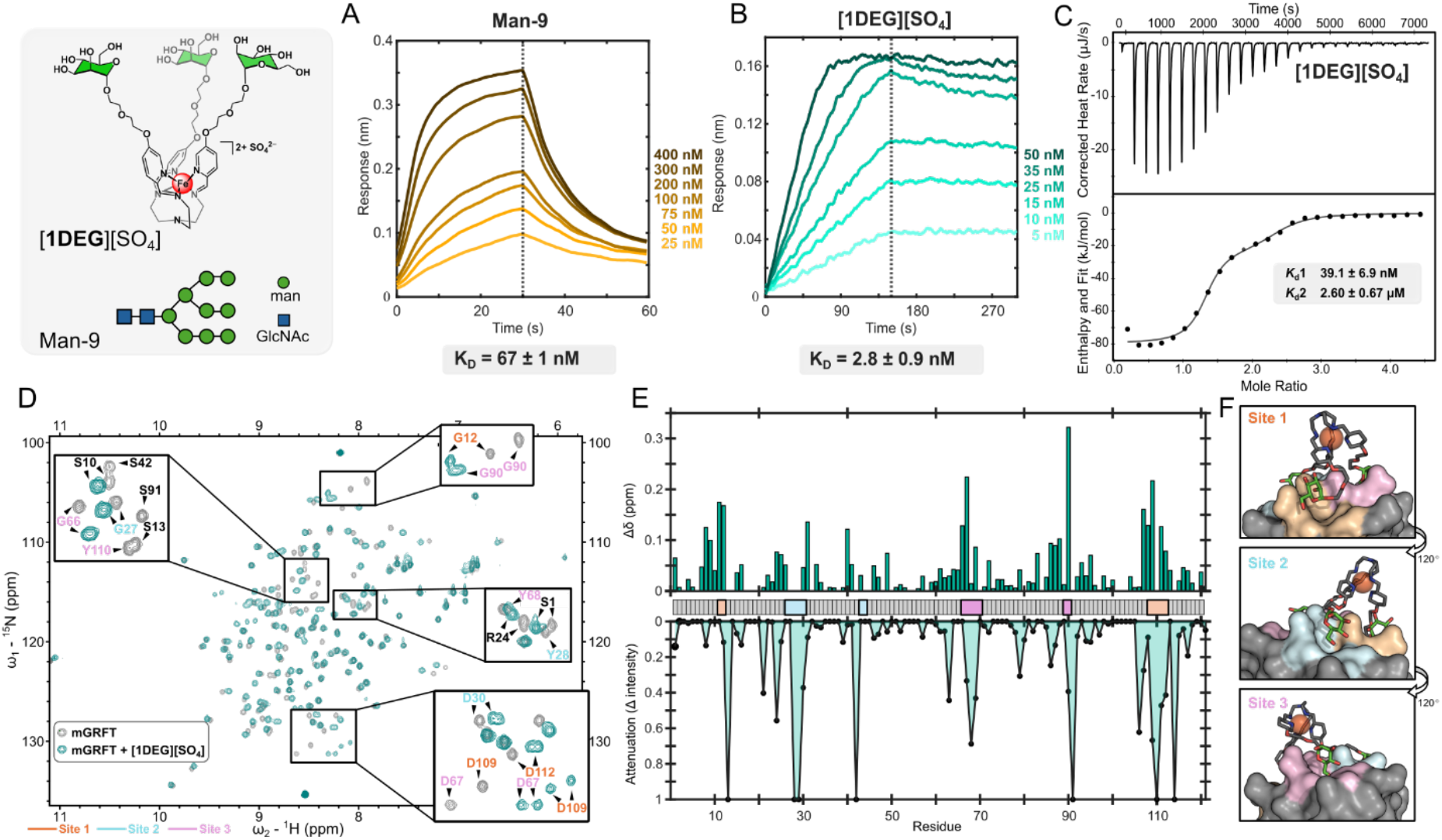
A) BLI trace with biotinylated Man-9 as the ligand and varying concentration of mGRFT as the analyte. B) BLI trace with mGRFT as the ligand with varying concentration of [**1DEG**]^2+^ as the analyte. C) ITC thermograms (top) and fitted binding isotherms (bottom) for the binding of [**1DEG**]^2+^ and mGRFT. D) 2D-^1^H ^15^N HSQC NMR spectra of mGRFT alone (grey) and mGRFT bound with [**1DEG**]^2+^ (teal). E) Chemical shift perturbations, top, and changes in signal intensity, bottom, as a function of amino acid residue. F) MOE docking of [**1DEG**]^2+^ onto mGRFT with the molecular surface colored based on binding site.

BLI measurements demonstrate that [**1DEG**]^2+^ binds mGRFT with low-nanomolar strength (2.8 ± 0.9 nM, Fig 3B), representing a nearly three-fold affinity gain over [**1**]^2+^. Remarkably, this single-EG extension drives the interaction beyond the affinity of mGRFT for its native ligand, Man-9. The BLI-derived binding kinetics suggest this 1,000-fold affinity enhancement is driven primarily by a substantially slower dissociation rate for [**1DEG**]^2+^ (k_d_ = 6.7 x 10^-4^ s^-1^), compared to the much faster off rate measured for [**1**]^2+^ (k_d_ = 1.6 x 10^-1^ s^-1^). These distinct kinetic profiles suggest that [**1DEG**]^2+^ and [**1**]^2+^ engage mGRFT differently, motivating the use of ITC to probe the binding behavior in more detail. Consistent with the two different binding profiles observed by BLI, ITC indicated that the interaction between the [**1DEG**]^2+^ analog and mGRFT is best described by a two-site model consisting of a high affinity interaction (*K*_d(avg)_ = 39.7 ± 6.9 nM) and a secondary, weaker interaction (*K*_d(avg)_ = 2.60 ± 0.67 μM), each with a stoichiometry of one (Fig 3C). These data contrast with the single-site model observed for the [**1**]^2+^, suggesting that extending the mannose tether allows the assembly to engage the same mGRFT monomer in two distinct binding events that each occur in a 1:1 stoichiometry. Consistent with this interpretation, docking studies (Fig. 3F) show that the extended arm length in [**1DEG**]^2+^ allows the complex to reach all three binding sites simultaneously, supporting the rationale behind our synthetic adjustment to drive avidity-driven binding through multisite engagement of a single molecule.

A 2D ^1^H-^15^N HSQC NMR spectrum of mGRFT bound to [**1DEG**]^2+^ provided further insight into this interaction. While the binding of [**1**]^2+^ with mGRFT primarily caused peak attenuations for Asp residues (D67, D109), the binding of [**1DEG**]^2+^ resulted in chemical shift changes and observable peak doubling for D67, D109, D30, and D112 (Fig 3D,E). These data suggest that there are two stabilized interactions in the [**1DEG**]^2+^/mGFRT bound state with slightly different chemical environments. Taken together with the BLI and ITC data, these observations support a binding model where [**1DEG**]^2+^ forms two strong contacts and a transient third interaction, which accounts for the slower dissociation and enhanced overall affinity when compared with the binding of the [**1**]^2+^ congener.

To further explore how these synthetic assemblies can be tuned to enhance mGRFT binding, we prepared [Fe(man_2_)_3_][SO_4_] ([**2**][SO_4_], Fig. 1B), in which each of the three terminal monosaccharides of [**1**]^2+^ is replaced with a 1,2-α-dimannose (mannobiose) unit to probe the effect of local di-mannose presentation. This modification yielded comparably tight binding to that of [**1DEG**]^2+^ as determined by BLI measurements, with a K_D_ of 3.2 ± 2.1 nM (Fig. 4A). Although the overall affinities are similar, differences in the association and dissociation rates suggest that [**2**]^2+^ may engage mGRFT through a different binding mechanism. [**1DEG**]^2+^ has a slower k_a_ (2.5 x 10^5^ M^-1^s^-1^) paired with a slow k_d_ (6.7 x 10^-4^ s^-1^) while [**2**]^2+^ has a faster k_a_ (1.0 x 10^6^ M^-^s^-^) and slightly quicker k_d_ (3.4 x 10^-3^ s^-1^) than [**1DEG**]^2+^ yet still slow in the context of ligand binding. Structural analysis by NMR spectroscopy revealed CSPs and peak attenuations for key binding site residues D67, D109, D30, and D112, reflecting behavior similar to that observed for the [**1DEG**]^2+^ complex. In addition, other resonances (G90, G66 and G12) exhibited a change in exchange profile from attenuation to chemical shift changes upon binding (Fig 4C,D), indicative of the formation of a stable complex. Similar to the [**1DEG**]^2+^ system, ITC analysis for the [**2**]^2+^/mGRFT interaction was fit to a two-site model consisting of a high-affinity interaction with *K*_d_ < 1 nM and a secondary interaction with *K*_d_ = 235.4 ±96.5 nM, indicating overall stronger binding for [**2**]^2+^ compared to that of [**1DEG**]^2+^(Fig. 4B). These differences in binding are consistent with the inherently stronger affinity of mGRFT for di-mannose compared to a single mannose residue,^35^ which likely reflects additional contacts formed with the second mannose unit.

**Figure 4:**
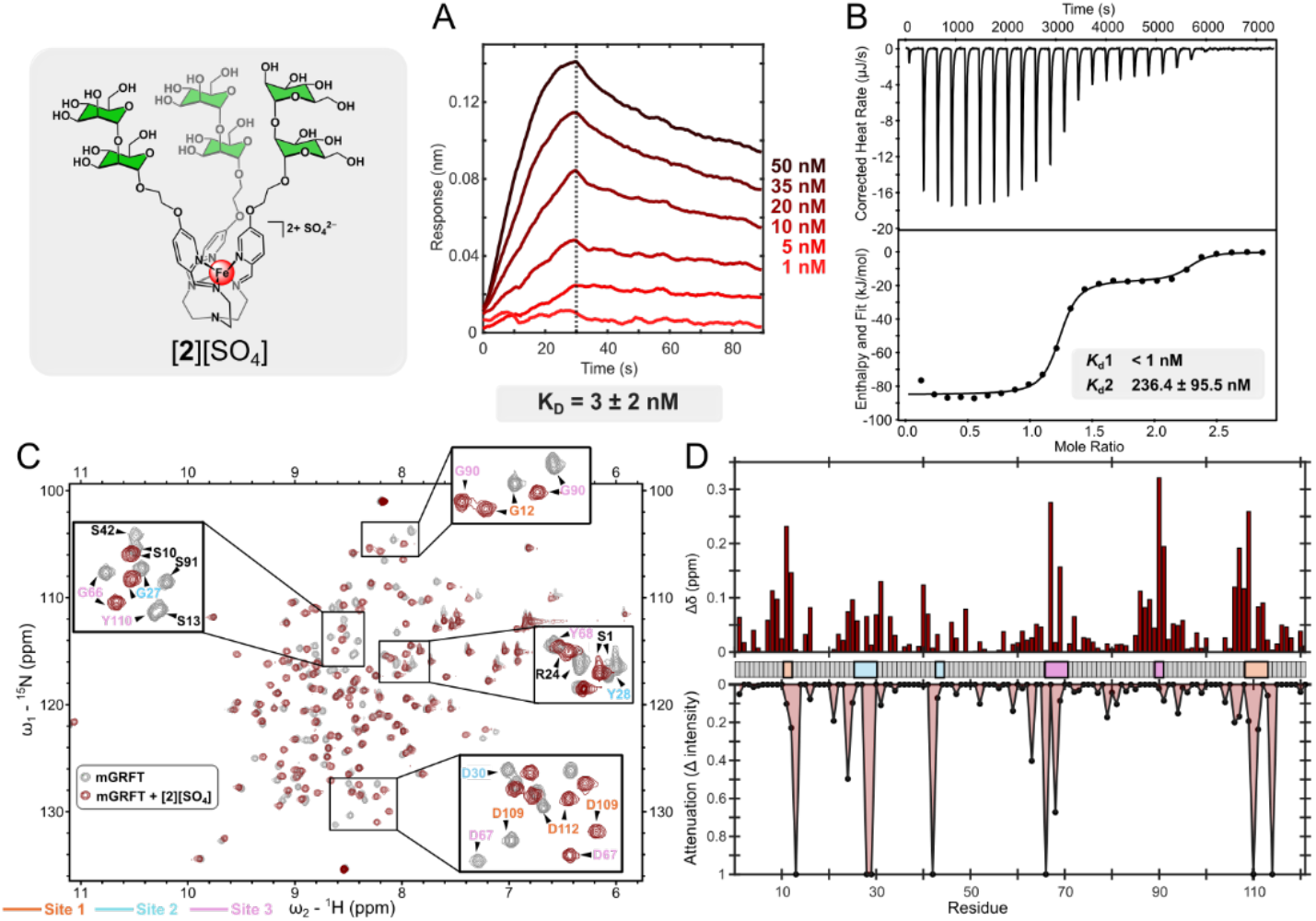
A) BLI trace with mGRFT as the ligand and varying concentration of [**2**]^2+^ as the analyte. B) ITC thermograms (top) and fitted binding isotherms (bottom) for the binding of [**2**]^2+^ and mGRFT. C) 2D ^1^H-^15^N HSQC NMR spectra of mGRFT alone (grey) and mGRFT bound with [**2**]^2+^ (maroon). D) Chemical shift perturbations, top, and changes in intensity, bottom, as a function of amino acid residue.

### [2DEG]^2+^ as a competitive inhibitor of high mannose glycan binding

Building on the significant binding enhancements observed for [**1DEG**]^2+^ and [**2**]^2+^, we next combined the diethylene glycol and di-mannose modifications to prepare the [**2DEG**]^2+^ analog. BLI analysis of the [**2DEG**]^2+^/mGRFT binding indicates an even stronger interaction for this dual-modified complex than what was observed for the [**1DEG**]^2+^ and [**2**]^2+^ systems. With an association rate of 1.4 x 10^6^ M^-1^s^-1^ and no measurable dissociation, [**2DEG**]^2+^ appears to combine the fast k_a_ of [**2**]^2+^ and the slow k_d_ of [**1DEG**]^2+^, resulting in an estimated K_D_ < 1 pM, which falls below the detection limit of BLI (Fig. 5A). We further characterized this binding using ITC. The resulting thermogram displayed extremely large exothermic responses during the initial injections, followed by a rapid transition to near-baseline heat release (Fig 5B). This profile is characteristic of ultra-high-affinity interactions, which preclude the determination of a dissociation constant by the technique, and in line with the magnitude of the binding strength observed by BLI. As anticipated, [**2DEG**]^2+^ exhibits NMR chemical shift and attenuation patterns similar to those observed for [**1DEG**]^2+^ and [**2**]^2+^ (Fig. 5 D,E), supporting the design strategy that combining the diethylene glycol extension with a di-mannose fragment is an effective method of generating an ultra-high affinity glycan mimetic.

**Figure 5:**
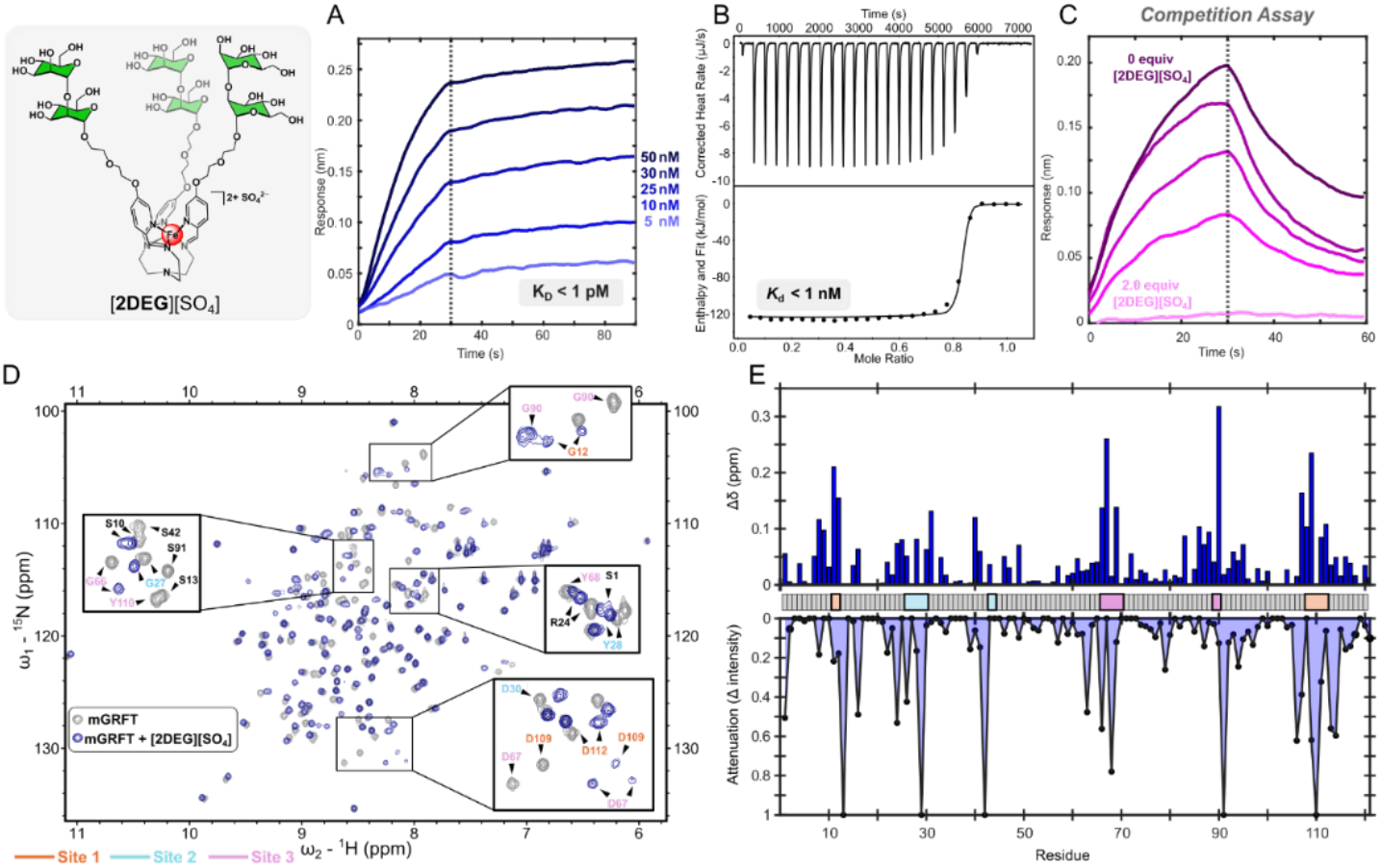
A) BLI trace with mGRFT as the ligand and varying concentration of [**2DEG**]^2+^ as the analyte. B) ITC thermograms (top) and fitted binding isotherms (bottom) for the binding of [**2DEG**]^2+^ and mGRFT. C) Competition assay with Man-9 as the ligand and mGRFT pre-treated with varying concentrations of [2DEG]2+ as the analyte. D) 2D-^1^H ^15^N HSQC of mGRFT alone (grey) and mGRFT bound with [**2DEG**]^2+^ (navy). (E) Chemical shift perturbations, top, and changes in signal intensity, bottom, as a function of amino acid residue.

To further probe this ultra-high-affinity interaction, we performed a BLI competition assay to determine whether the [**2DEG**]^2+^ complex could outcompete mGRFT’s natural ligand, Man-9. Biotinylated Man-9 was immobilized on the BLI biosensor tip, which was then dipped into solutions containing mGRFT that was pre-incubated with varying concentrations of [**2DEG**]^2+^ A concentration-dependent decrease in signal during the association phase was observed, with 0.5 molar equivalents of [**2DEG**]^2+^ reducing the response by approximately 50% and two molar equivalents nearly eliminating the response entirely (Fig. 5C). These results indicate that mGRFT preferentially binds to [**2DEG**]^2+^, enabling the synthetic complex to outcompete its native Man-9 ligand. This result demonstrates that the tailored sugar arrangement and preorganized architecture of the [**2DEG**]^2+^ framework can produce a binding interaction that surpasses the strength of the natural high-mannose presentation recognized by GRFT in nature. More broadly, this finding highlights the power of rational synthetic design to generate glycomimetics capable of outperforming their biological counterparts.

## Conclusion

This work demonstrates that iminopyridine-supported Fe(II) glycan assemblies offer a highly tunable synthetic platform for generating precise mimetics of N-linked high-mannose glycans. When mannose residues are displayed from the iminopyridine ligand framework *via* a single ethylene glycol linker, the resulting [**1**]^2+^ assembly mimics the binding of a Man-5 glycan as evidenced by the direct comparison of NMR signatures for both systems. Extension of the linker arms by an additional ethylene glycol unit result in a 1000-fold enhancement in mGRFT binding, which is driven primarily by a significantly slowed dissociation rate. The combined BLI, ITC, and NMR analyses reveal that this enhancement arises from a binding mode in which [**1DEG**]^2+^ engages mGRFT through a dominant high-affinity contact accompanied by a transient secondary interaction, which together surpasses both the affinity and kinetic stability of the parent [**1**]^2+^ complex. Addition of a second appended mannose residue to the molecular architecture resulted in a binding profile similar to that observed for [**1DEG**]^2+^, with comparable affinity and kinetic characteristics. Combining the diethylene glycol extension with a second appended mannose residue produced the [**2DEG**]^2+^ assembly, which engages in an ultra-high-affinity interaction with mGRFT that surpasses the binding strength of the parent complexes and can outcompete mGRFT’s natural Man-9 ligand. Uniquely, the tunable and combinatorial nature of these Fe(II)-anchored glycan assemblies provides precise control over terminal mannose presentation to enable the design of high-affinity glycomimetics capable of outcompeting native biological ligands. This platform provides a versatile strategy for modeling HMGs and opens avenues for the development of competitive inhibitors targeting medically-relevant protein-glycan interactions.

## Methods

### Protein Purification

Monomeric griffithsin (mGRFT) was purified from *E. coli* cells using a Cytiva HisTrap HP column and a HiTrap Q HP column. Briefly, *E. coli* BL21(DE3) were transformed with a pET28b+ plasmid containing a N-terminal hexa-his tag followed by a TEV cleavage site and the sequence for monomeric Grifithsin.^33^ Cells were grown to an OD600 of 0.6 and induced with 1 mM IPTG at 18 °C for 16 h. Cells were harvested by centrifugation at 3000 x g, the supernatant was discarded, and the pellet was resuspended into wash buffer (1x PBS, 200mM NaCl, 20mM Imidazole pH 7.4). Cells were lysed via sonication (amplitude 60, 30s on with 45s rest for 6 min). The lysate was centrifuged at 18,000 x g for 45 minutes and the supernatant was filtered through a 0.2 µM filter. Subsequently, the clarified lysate was loaded on a HisTrap HP column and eluted with an imidazole gradient (5-100% imidazole over 10 CVs). The NiNTA eluate was dialyzed into 20mM Tris pH 7.5 then loaded onto a HiTrap Q HP column and eluted with a salt gradient (0-1M NaCl over 20 CVs). The purity and identity was established via SDS-PAGE.

### NMR spectroscopy

All protein NMR spectra were collected at 298 K in 1 x PBS buffer pH 7.4 with 10% D_2_O using a Bruker Avance Neo 800 MHz spectrometer equipped with a triple resonance TXO cryoprobe. 2D ^1^H-^15^N HSQC spectra were collected using 125 μM of His-TEV mGRFT with varying concentrations of Fe(II) glycomimetics. All spectra were processed in Topspin and visualized in POKY.^41^ Assignments were transferred from BMRB entry #18585.^36^ Raw peak intensities were determined via integration with a gaussian fit then normalized with in the spectra to a standard peak. Change in intensity was determined using the formula 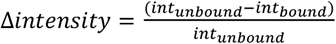 CSPs were calculated after internally referencing to Trimethylsilylpropanoic acid.

### Biolayer Interferometry

All BLI experiments were performed at 25 °C on an Octet R2 instrument using SA or NiNTA biosensors. Ligand was loaded onto the biosensors followed by a baseline then association and dissociation phase. Biosensors were regenerated and this process was repeated for varying concentrations of analyte. Data was processed in Octet analysis studio 13.0. The samples were double referenced, first to a reference sensor then to a well containing no analyte. The trace responses were aligned to the average of the final 5s of the baseline and inter-step corrected to the dissociation step. A Savitzky-Golay filter was applied to reduce high frequency noise. The association and dissociation phases were fit globally to a 1:1 single exponential binding model. All conditions were run in triplicate.

### Molecular Docking

Docking was performed in MOE using mGRFT (PDB: 3LL2) as the receptor and [**1DEG**]^2+^ or **[1]**^**2+**^ as the ligands. Placement was based on triangle matching with London dG scoring while refinement was performed with a rigid receptor fit with GBVI/WSA dG. 100 poses were sampled and the top 10 were retained. Docking results were visualized using PYMOL.^42^

## Acknowledgements

A.J.G. acknowledges support from R00GM145970 and U54CA272220. J.M.S is supported by the National Institute of General Medical Sciences of the National Institutes of Health through a Maximizing Investigators’ Research Award (MIRA) (R35GM160296) and through UCSD startup funds. The authors acknowledge the use of facilities and instrumentation supported by the National Science Foundation through the University of California San Diego Materials Research Science (MRSEC) and Engineering Center (DMR-2011924). The authors thank Dr. Xuemei Huang and Dr. Anthony Mrse for maintaining the NMR spectrometers and Dr. Elizabeth Komives for discussions of BLI and ITC data.

